# Unexpectedly high mutation rate of a deep-sea hyperthermophilic anaerobic archaeon

**DOI:** 10.1101/2020.09.09.287623

**Authors:** Jiahao Gu, Xiaojun Wang, Xiaopan Ma, Ying Sun, Xiang Xiao, Haiwei Luo

## Abstract

Deep-sea hydrothermal vents resemble the early Earth, and thus the dominant *Thermococcaceae* inhabitants, which occupy an evolutionarily basal position of the archaeal tree and take an obligate anaerobic hyperthermophilic free-living lifestyle, are likely excellent models to study the evolution of early life. Here, we determined that unbiased mutation rate of a representative species, *Thermococcus eurythermalis*, exceeded that of all known free-living prokaryotes by 1-2 orders of magnitude, and thus rejected the long-standing hypothesis that low mutation rates were selectively favored in hyperthermophiles. We further sequenced multiple and diverse isolates of this species and calculated that *T. eurythermalis* has a lower effective population size than other free-living prokaryotes by 1-2 orders of magnitude. These data collectively indicate that the high mutation rate of this species is not selectively favored but instead driven by random genetic drift. The availability of these unusual data also helps explore mechanisms underlying microbial genome size evolution. We showed that genome size is negatively correlated with mutation rate and positively correlated with effective population size across 30 bacterial and archaeal lineages, suggesting that increased mutation rate and random genetic drift are likely two important mechanisms driving microbial genome reduction. Future determinations of the unbiased mutation rate of more representative lineages with highly reduced genomes such as *Prochlorococcus* and *Pelagibacterales* that dominate marine microbial communities are essential to test these hypotheses.

---

One theory for the origin of life is that the last universal common ancestor was an anaerobic hyperthermophilic organism inhabiting the deep sea hydrothermal vents, as these environments display a few characteristics paralleling the early Earth [1]. While hydrothermal vents vary with chemical parameters, they all share a high temperature zone near the black chimney with anaerobic fluid from it. In the past decades, great efforts were made to understand the metabolic strategies deep-sea hyperthermophiles use to conserve energy and cope with physicochemical stresses, and to appreciate the molecular mechanisms leading to the stabilization of nucleic acids and proteins at exceedingly high temperatures [2, 3]. However, little is known whether they have a low or high intrinsic (i.e., not selected by environmental pressure) rate to change their genetic background information and whether this intrinsic potential itself is a result of selection shaped by these unique habitats.

A previous population genomic analysis showed that protein sequences are under greater functional constraints in thermophiles than in mesophiles, suggesting that mutations are functionally more deleterious in thermophiles than in mesophiles [4]. This explanation is also supported by experimental assays showing nearly neutral mutations in temperate conditions become strongly deleterious at high temperature [5]. Furthermore, fluctuation tests on a hyperthermophilic archeaon *Sulfolobus acidocaldarius* [6] and a hyperthermophilic bacterium *Thermus thermophilus* [7] consistently showed that hyperthermophiles have much lower mutation rate compared to mesophiles. This appears to support the hypothesis that selection favors high replication fidelity at high temperature [5].

Nevertheless, mutation rates measured using fluctuation experiments based on reporter loci are known to be biased, since the mutation rate of the organism is extrapolated from a few specific nonsynonymous mutations enabling survival in an appropriate selective medium, which renders the results susceptible to uncertainties associated with the representativeness of these loci and to inaccuracies of the assumptions made in extrapolation methods [8–10]. These limitations are avoided by the mutation accumulation (MA) experiment followed by whole-genome sequencing (WGS) of derived lines. In the MA part, multiple independent MA lines initiated from a single progenitor cell each regularly pass through a single-cell bottleneck, usually by transferring on solid medium. As the effective population size (*N*_*e*_) becomes one, selection is unable to eliminate all but the lethal mutations, rendering the MA/WGS an approximately unbiased method to measure the spontaneous mutation rate [11].

Members of the free-living anaerobic hyperthermophilic archaeal family *Thermococcaceae* are among the dominant microbial lineages in the black-smoker chimney at Guaymas Basin [12] and other deep sea hydrothermal vents [13, 14]. This family only contains three genera: *Thermococcus, Pyrococcus* and *Palaeococcus*. In this study, the MA/WGS procedure was applied to determine the unbiased spontaneous mutation rate of a representative member *Thermococcus eurythermalis* A501, a conditional pizeophilic archaeon which can grow equally well from 0.1 MPa to 30 MPa at 85°C [15, 16]. The MA lines were propagated at this optimal temperature on plates with gelrite which tolerates high temperature, and the experiment was performed under normal air pressure and in strictly anaerobic condition (Fig. 1A-D). To the best of our knowledge, this is the first report of unbiased mutation rate of a hyperthermophile and an obligate anaerobe.

**Figure 1.**
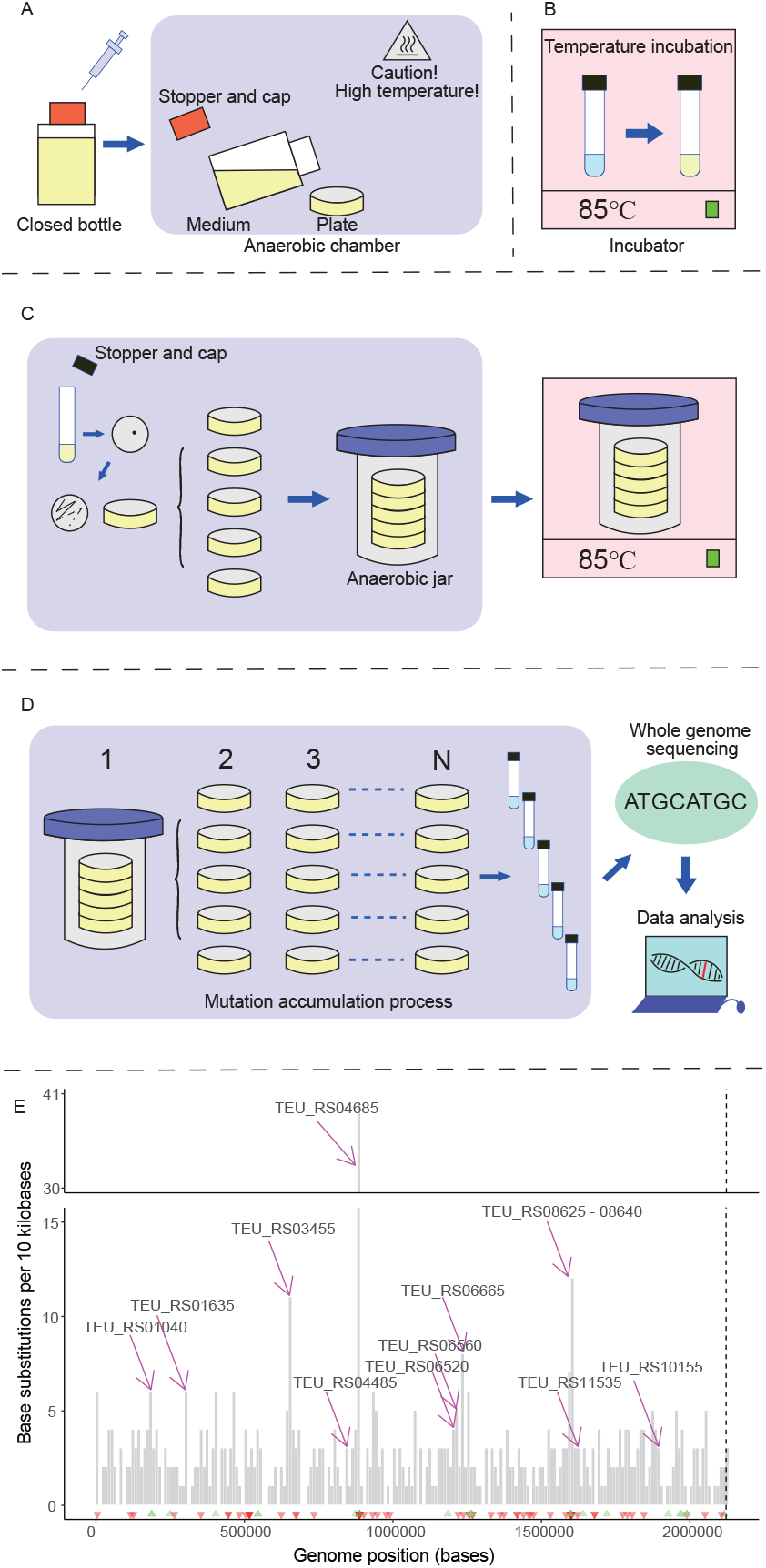
Experimental determination of the unbiased mutation rate of the *Thermococcus eurythermalis* A501 is challenging because this archaeon has unusual physiology (i.e., obligate anaerobic and obligate hyperthermophilic). **(A)** The preparation of anaerobic high temperature tolerant gelrite plate. After sterilization and polysulfide addition via syringe, the plates are made in an anaerobic chamber. **(B)** The incubation of the strain *T. eurythermalis* A501 at 85°C in liquid medium. **(C)** The initiation of mutation accumulation (MA) by spreading cells from a single founding colony to 100 lines. Plates are placed in an anaerobic jar for incubation in strictly anaerobic condition at 85°C. **(D)** The MA process followed by whole genome sequencing and data analysis. Single colony of each line is transferred to a new plate for N times (here N=20). **(E)** Base-substitution mutations and insertion/deletion mutations across the whole genome of *T. eurythermalis*. The dashed vertical line separates the chromosome and plasmid. The height of each bar represents the number of base-substitution mutations across all MA lines within 10 kbp window. Green and red triangles denote insertion and deletion, respectively. The locus tags of the 14 genes with statistical enrichment of mutations are shown.

Our MA experiment allowed accumulation of mutations over 314 cell divisions (after correcting the death rate (Table S1) [17]) in 100 independent lines initiated from a single founder colony and passed through a single cell bottleneck every day. By sequencing genomes of 96 survived lines at the end of the MA experiment, we identified 544 base-substitution mutations over these lines (Table S2), which translates to an average mutation rate (µ) of 85.01×10^−10^ per cell division per nucleotide site (see Supplementary information). The ratio of accumulated nonsynonymous to synonymous mutations (371 vs 107) did not differ from the ratio of nonsynonymous to synonymous sites (1,485,280 vs 403,070) in the A501 genome (*χ*^*2*^ test; *p*>0.05). Likewise, there was no difference of the accumulated mutations between intergenic (65) and protein-coding sites (478) (*χ*^*2*^ test; *p*>0.05). These are evidence for minimal selective elimination of deleterious mutations during the MA process. In general, the mutations were randomly distributed along the chromosome and the plasmid, though 86 mutations fell into 14 genes which showed significant enrichment of mutations (bootstrap test; *p*<0.05 for each gene) and 52 out of the 86 mutations were found in five genes (TEU_RS04685 and TEU_RS08625-08640 gene cluster) (Fig. 1E, Table S3). A majority of these mutations led to stop codons and thus may have inactivated these genes (38 out of 71 in the former gene and 33 out of 43 in the latter gene cluster). The phenomenon of mutation clustering is not unique to this organism; it was reported in another MA study with the yeast *Schizosaccharomyces pombe*, and these genomic regions may represent either mutational hotspots or that mutations confer selective advantages under experimental conditions [18]. The TEU_RS04685 encodes the beta subunit of the sodium ion-translocating decarboxylase which is an auxiliary pathway for ATP synthesis by generating sodium motive force via decarboxylation [19], and the TEU_RS08625-08640 encodes a putative nucleoside ABC transporter. These genes appear to be important for energy conservation in the highly fluctuating deep-sea hydrothermal fluids. Under the culture conditions in which peptides and amino acids were stably and sufficiently supplied (see the TRM medium recipe in Supplementary information), however, these genes may be dispensable because peptides and amino acids are the preferred carbon and energy sources for *T. eurythermalis* [15]. On the other hand, some of these genes (e.g., TEU_RS08625) were shown to be upregulated under alkaline stress [16], and thus may be similarly induced under the culture condition in which pH is elevated compared to the vents. Besides, the laboratory condition differed from the vents in a number of other physicochemical features including hydrostatic pressure (0.1 MPa during the MA process versus 20 MPa *in situ*), temperature and salinity, which likely imposed additional selective pressures on the mutation accumulation processes. Taken together, deleting these genes were likely translated to a net fitness gain and were thus driven by selection. Removing these mutations led to a spontaneous mutation rate of 71.57×10^−10^ per cell division per site for *T. eurythermalis* A501. After removing the mutations in these 14 genes, both the accumulated mutations at nonsynonymous sites (288) relative to those (104) at synonymous sites (*χ*^*2*^ test; *p*=0.014) and the accumulated mutations at intergenic regions (65) relative to protein-coding regions (392) (*χ*^*2*^ test; *p*=0.013) showed marginally significant differences.

To date, over 20 phylogenetically diverse free-living bacterial species and two archaeal species isolated from various environments have been assayed with MA/WGS, and their mutation rates vary from 0.79×10^−10^ to 97.80×10^−10^ per cell division per site [20]. The only prokaryote that displays a mutation rate (97.80×10^−10^ per cell division per site) comparable to A501 is *Mesoplasma florum* L1 [21], a host-dependent wall-less bacterium with highly reduced genome (∼700 genes). Our PCR validation of randomly chosen 20 base-substitution mutations from two MA lines displaying highest mutation rates and of all nine insertion-deletion (INDEL) mutations involving >10 bp changes across all lines (Table S2) indicates that the calculated high mutation rate did not result from false bioinformatics predictions.

The extremely high mutation rate of *T. eurythermalis* is unexpected. One potential explanation in line with the “mutator theory” [22–24] is that high mutation rate may allow the organisms to gain beneficial mutations more rapidly and thus is selectively favored in deep-sea hydrothermal vents where physicochemical parameters are highly fluctuating. Alternatively, high mutation rate is the result of random genetic drift according to the “drift-barrier model” [21]. In this model, increased mutation rates are associated with increased load of deleterious mutations, so natural selection favors lower mutation rates. On the other hand, increased improvements of replication fidelity come at an increased cost of investments in DNA repair activities. Therefore, natural selection pushes the replication fidelity to a level that is set by genetic drift, and further improvements are expected to reduce the fitness advantages [11, 21]. These two explanations for the high mutation rate of *T. eurythermalis* are mutually exclusive, and resolving them requires the calculation of the power of genetic drift, which is inversely proportional to *N*_*e*_.

A common way to calculate *N*_*e*_ for a prokaryotic population is derived from the equation *π*_*S*_ =2×*N*_*e*_×µ, where *π*_*S*_ represents the nucleotide diversity at synonymous (silent) sites among randomly sampled members of a panmictic population [25]. We therefore sequenced genomes of another eight *T. eurythermalis* isolates available in our culture collections. Like *T. eurythermalis* A501, these additional isolates were collected from the same cruise but varying at the water depth from 1,987 m to 2,009 m at Guaymas Basin. They differ by only up to 0.135% in the 16S rRNA gene sequence and share a minimum whole-genome average nucleotide identity (ANI) of 95.39% (Table S4), and thus fall within an operationally defined prokaryotic species typically delineated at 95% ANI [26]. Population structure analysis with PopCOGenT [27] showed that these isolates formed a panmictic population and that two of them were repetitive as a result of clonal descent (see Supplementary information). Using the median value of *π*_*S*_ =0.083 across 1,628 single-copy orthologous genes shared by the seven non-repetitive genomes, we calculated the *N*_*e*_ of *T. eurythermalis* to be 5.83×10^6^.

Next, we collected the unbiased mutation rate of other prokaryotic species determined with the MA/WGS strategy from the literature [11, 28–30]. While the *N*_*e*_ data were also provided from those studies, the isolates used to calculate the *N*_*e*_ were identified based on their membership of either an operationally defined species (e.g., ANI at 95% cutoff) or a phenotypically characterized species (e.g., many pathogens), which often create a bias in calculating *N*_*e*_ [25]. We therefore again employed PopCOGenT to delineate panmictic populations from those datasets and re-calculated *N*_*e*_ accordingly. There was a significant negative linear relationship between µ and *N*_*e*_ on a logarithmic scale (dashed gray line in Fig. 2A [r^2^ = 0.83, slope = -0.85, s.e.m. = 0.09, *p*<0.001]) according to a generalized linear model (GLM) regression. This relationship cannot be explained by shared ancestry, as confirmed by phylogenetic generalized least square (PGLS) regression analysis (solid blue line in Fig. 2A [r^2^ = 0.81, slope = -0.81, s.e.m. = 0.09, *p*<0.001]). The nice fit of *T. eurythermalis* to the regression line validated the drift-barrier hypothesis. This is evidence that the high mutation rate of *T. eurythermalis* is driven by genetic drift rather than by natural selection.

**Figure 2.**
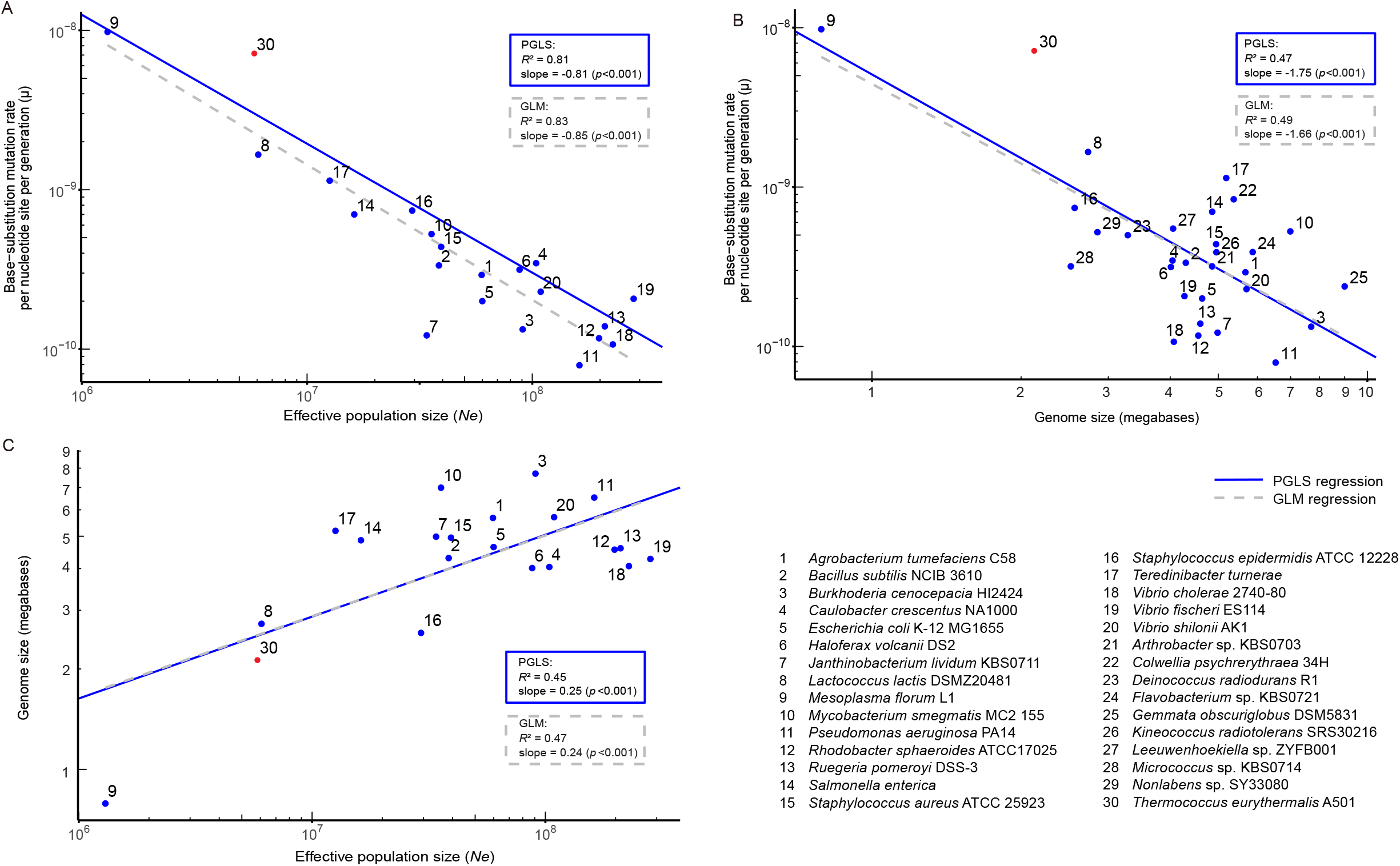
The scaling relationship involving the base-substitution mutation rate per cell division per site (µ), the estimated effective population size (*N*_*e*_), and genome size across 28 bacterial and two archaeal species. All three traits’ values were logarithmically transformed. The mutation rates of these species are all determined with the mutation accumulation experiment followed by whole genome sequencing of the mutant lines. The mutation rate of species numbered 1-29 (blue) is collected from literature and that of the species 30 (red) is determined in the present study. Among the numbered species shown in the figure, the species #6 *Haloferax volcanii* is facultative anaerobic halophilic archaeon, and the species #30 is an obligate anaerobic hyperthermophilic archaeon. **(A)** The scaling relationship between µ and *N*_*e*_. **(B)** The scaling relationship between µ and genome size. **(C)** The scaling relationship between genome size and *N*_*e*_. Numbered data points 21-29 are not shown in (A) and (C) because of the lack of population dataset for estimation of *N*_*e*_. The dashed gray lines and blue lines represent the generalized linear model (GLM) regression and the phylogenetic generalized least square (PGLS) regression, respectively. The Bonferroni adjusted outlier test for the GLM regression show that #7 *Janthinobacterium lividum* is an outlier in the scaling relationship between µ and *N*_*e*_, and #9 *Mesoplasma florum* is an outlier in the scaling relationship between genome size and *N*_*e*_. No outlier was identified in the PGLS regression results.

As stated in the drift-barrier theory, high mutation rate is associated with a high load of deleterious mutations. In the absence of back mutations, recombination becomes an essential mechanism in eliminating deleterious mutations [31]. In support of this argument, the ClonalFrameML analysis [32] shows that members of the *T. eurythermalis* population recombine frequently, with a high ratio of the frequency of recombination to mutation (ρ/θ=0.59) and a high ratio of the effect of recombination to mutation (r/m=5.76). In fact, efficient DNA incorporation to *Thermococcaceae* genomes from external sources has been well documented experimentally [33, 34]. A second potentially important mechanism facilitating *T. eurythermalis* adaptation at high temperature is strong purifying selection at the protein sequence level, as protein sequences in thermophiles are generally subjected to stronger functional constraints compared to those in mesophiles [4, 35].

Our result of the exceptionally high mutation rate of a free-living archaeon is a significant addition to the available collection of the MA/WGS data (Table S5), in which prokaryotic organisms with very high mutation rate have only been known for a host-dependent bacterium (*Mesoplasma florum* L1) with unusual biology (e.g., cell wall lacking). The availability of these two deeply branching (one archaeal versus the other bacterial) organisms adopting opposite lifestyles (one free-living versus the other host-restricted; one hyperthermophilic versus the other mesophilic; one obligate anaerobic versus the other facultative anaerobe), along with other phylogenetically and ecologically diverse prokaryotic organisms displaying low and intermediate mutation rates, provides an opportunity to help illustrate mechanisms potentially driving genome size evolution across prokaryotes. We found a negative linear relationship (dashed gray line in Fig. 2B [r^2^ = 0.49, slope = -1.66, s.e.m. = 0.32, *p*<0.001]) between genome size and base-substitution mutation rate, which is consistent with the hypothesis that increased mutation rate drives microbial genome reduction. We also showed a positive linear relationship (dashed gray line in Fig. 2C [r^2^ = 0.47, slope = 0.24, s.e.m. = 0.06, *p*<0.001]) between genome size and *N*_*e*_, which suggests that random genetic drift drives genome reduction across prokaryotes. These correlations remain robust when the data were analyzed as phylogenetically independent contrasts (blue solid lines in Fig. 2B [r^2^ = 0.47, slope = -1.75, s.e.m. = 0.34, *p*<0.001] and in Fig. 2C [r^2^ = 0.45, slope = 0.25, s.e.m. = 0.06, *p*<0.001]). Our results are consistent with recent studies which employed mathematical modeling and/or comparative sequence analyses to show random genetic drift [36] and increased mutation rate [37] driving genome reduction across diverse bacterial lineages including both free-living and host-dependent bacteria. One benefit of the present study is that it directly measures µ and *N*_*e*_, as compared to those recent advances which relied on proxies for these metrics (e.g., using the ratio of nonsynonymous substitution rate to synonymous substitution rate to represent *N*_*e*_) to infer mechanisms of genome reduction.

Despite this advantage, there are important caveats to our conclusions related to the mechanisms of genome reduction. The correlation analyses performed here are inspired by Lynch and colleagues’ work, who had great success explaining eukaryotic genome expansion with genetic drift [11, 38]. However, there are a few key differences of genomic features between prokaryotes and eukaryotes, which makes it more difficult to explain the correlation observed in prokaryotes. Importantly, genome sizes of eukaryotes can vary over several orders of magnitude, whereas those of free-living prokaryotes differ by only an order of magnitude [11], so there is much less variability to explain in prokaryotes. Moreover, eukaryotic genomes experience dramatic expansions of transposable elements which are often considered as genomic parasites, whereas prokaryotic genomes including those large ones are usually depleted with transposable elements and their genome size variations are largely driven by gene content [39]. Aside from these conceptual difficulties, the plots (Fig. 2B & 2C) are poorly populated with typical free-living species carrying small genomes such as the *Prochlorococcus* (mostly 1.6-1.8 Mb) and *Pelagibacterales* (1.3-1.5 Mb), which dominate the photosynthetic and heterotrophic microbial communities, respectively, in the ocean [40]. It has been generally postulated that bacterial species in these lineages have very large *N*_*e*_ [39–41], though there has been little direct evidence for it [42, 43]. If confirmed through the measurement of the unbiased mutation rate (µ) followed by the calculation of *N*_*e*_ based on µ, it might compromise the linear relationship between genome size and *N*_*e*_ observed here (Fig. 2C). It is also not necessarily appropriate to translate correlations to causal relationships. For example, the correlation between increased mutation rates and decreased genome sizes (Fig. 2B) does not necessarily mean that increased mutation rate drives genome reduction. This is because high mutation rates are observed in species with small *N*_*e*_. Given that deletion bias is commonly found in prokaryotes [44, 45], genome reduction can be easily explained by increased fixation of deletional mutations in species with smaller *N*_*e*_. High mutation rates in these species are simply the result of random genetic drift as explained by the drift-barrier theory, and they may have a limited role in driving genome reduction.

Whereas our analysis based on the available data did not support natural selection as a universal mechanism driving genome reduction across prokaryotes (Fig. 2B&C), it does not mean that selection has no role in genome reduction of a particular taxon. In the case of thermophiles, proponents for selection acting to reduce genomes explained that genome size, due to its positive correlation with cell volume, may be an indirect target of selection which strongly favors smaller cell volume [35]. The underlying principle is that high temperature requires cells to increase the lipid content and change the lipid composition of the cell membranes, which consumes a large part of the cellular energy, and thus lower cell volume is selectively favored at high temperature [35]. Our calculations of a relatively small *N*_*e*_ in *T. eurythermalis* does not necessarily contradict with this selective argument, given that the fitness gained by decreasing cell volume and thus reducing genome size is large enough to overcome the power of random genetic drift. On the other hand, our data strongly indicate that neutral forces dictate the genome evolution of *T. eurythermalis*, and they are not negligible with regard to its genome reduction process. The significantly more deletion over insertion events (t test; 95 versus 37 events with *p*<0.001 and 48 versus 20 events with *p*<0.05 before and after removing the 14 genes enriched in mutations, respectively) and the significantly more nucleotides involved in deletions over insertions (t test; 433 versus 138 bp with *p*<0.05 and 386 versus 121 bp with *p*<0.001 before and after removing the 14 genes enriched in mutations, respectively) suggest that the deletion bias, combined with increased chance fixation of deletion mutants due to low *N*_*e*_, is a potentially important neutral mechanism giving rise to the small genomes of *T. eurythermalis* (2.12 Mbp).

The globally distributed deep-sea hydrothermal vents are microbe-driven ecosystems, with no known macroorganisms surviving at the vent fluids. Sample collections, microbial isolations, and laboratory propagations of mutation lines at hot and anoxic conditions are challenging. In the present study, we determined that *T. eurythermalis*, and perhaps *Thermococcaceae* in general, has a highly increased mutation rate and a highly decreased effective population size compared to all other known free-living prokaryotic lineages. While it remains to be tested whether this is a common feature among the vents’ microbes, the present study nevertheless opens a new avenue for investigating the hyperthemophile ecology and evolution in the deep sea.

## Acknowledgements

This research is supported by the National Key R&D Program of China (2018YFC0309800), National Nature of Science China (NSFC 41530967), China Ocean Mineral Resources R & D Association DY125-22-04, and the Hong Kong Research Grants Council Area of Excellence Scheme (AoE/M-403/16).

## Author Contributions

H.L. conceptualized the work and strategy, directed the bioinformatics analyses, interpreted the data, and wrote the main manuscript. X.X. set up the experimental platform for deep sea hyperthermophile studies, directed the experimental analyses and related writing, co-interpreted the data, provided comments to the main manuscript, and acquired the strains. J.G. performed all the experiments with contributions from X.M, drafted the related methods in supplementary information and Figure 1. X.W. performed all the bioinformatics analyses, co-interpreted the related results, drafted the related methods in supplementary information, Figure 2 and all supplemental tables. Y.S. contributed the bioinformatics tools for mutation detection and mutation rate calculation.

## Data availability

All the datasets generated, analyzed, and presented in the current study are available in the Supplementary Information. Genomic sequences of the eight *Thermococcus eurythermalis* strains are available at the JGI IMG under the GOLD study id Gs0142375. Raw reads of the eight strains are available at the NCBI SRA under the accession number PRJNA679699.

## Code availability

The custom scripts used in this study have been deposited in the online repository (https://github.com/Xiaojun928/Mutation-calling).

