## Supplementary information for "Unexpectedly high mutation rate of a deep-sea hyperthermophilic anaerobic archaeon"

**This PDF file includes:**

Supplementary Materials and Methods

References

### Materials and Methods

*Sampling, cultivation, and genome sequencing of Thermococcus eurythermalis isolates*

Nine *Thermococcus eurythermalis* strains (Table S3) were isolated from samples of Guaymas Basin hydrothermal vents in the cruise number AT 15–55, during 7-17 November 2009 [1]. Briefly, samples were stored in the Hungate anaerobic tubes and kept at 4℃. Then the samples were enriched at 85℃ or 95℃ using *Thermococcales* Rich Medium (TRM). One liter of TRM contains 3.3g pipes disodium salt, 23g NaCl, 5g $MgCl_2 \cdot 6H_2O$, 0.7g KCl, 0.5g $(NH_4)_2SO_4$, 1mL $K_2HPO_4$ 5%, 1mL $KH_2PO_4$ 5%, 1mL $CaCl_2 \cdot 2H_2O$ 2%, 0.05g NaBr, 0.01g $SrCl_2 \cdot 6H_2O$, 1mL $Na_2WO_4$ 10mM, 1mL $FeCl_3$ 25mM, 1g yeast extract, 4g tryptone and 1mg resazurin [2]. The medium was adjusted to pH 7.0, autoclaved and reduced with 0.5g sodium sulphide before use. Next, enrichment cultures were inoculated on the solid medium prepared with hungate roll-tube technique and incubated at 85℃ or 95℃ under atmosphere pressure. Single colonies were transferred into new TRM medium and purified using roll-tube technique for 3 times and stocks were kept at -80℃. More details of sampling and isolation can be found in a previous paper [1]. Among these isolates, the complete genome of the type strain A501 (GCA_000769655.1) was downloaded from the NCBI GenBank database [3], and the rest eight strains were sequenced in the present study. To get enrichment of these eight strains, stocks kept in -80℃ were inoculated into 50 mL anaerobic TRM medium in the serum bottle and cultured in the incubator in 85℃. The liquid medium was supplemented with sulfur and $Na_2S \cdot 9H_2O$. After enrichment, the cells were collected using centrifuge (12,000 rpm, 10min). Genomic DNA of each isolate was extracted using the Magen Hipure

Soil DNA Kit and was sequenced using the Illumina Hiseq platform with 2×150 bp paired-end. Raw reads were first processed by Trimmomatic 0.32 [4] to remove adaptors and trim bases of low quality. The draft genome of each isolate was assembled with quality reads using SPAdes v3.10.1 [5] with default parameters.

*Mutation accumulation experiment*

For culture propagation under high temperature, anaerobic high-temperature-tolerant plates were made every day before the transfer. Plates were made using anaerobic *Thermococcus* Rich Medium [2] (TRM) with gelrite (15g liter$^{-1}$). After sterilization, 1.5 mL of a polysulfide solution [6] was added per liter of medium using syringe to make sure a strictly anaerobic condition. The medium was transferred into an anaerobic chamber (COY, Vinyl Anaerobic Chamber) immediately, preventing it from cooling. This is because gelrite used for making plates becomes solidified soon after it become cooler. Plates were made in the chamber.

The mutation accumulation (MA) experiment started from a single founder colony of *Thermococcus eurythermalis* A501. It was transferred to new plates to form 100 independent lines. Plates were put into an anaerobic jar (GeneScience), which were together moved to an incubator. After incubation at 85°C under normal air pressure (optimal growth pressure from 0.1-30 MPa) for one day, the jar was transferred back into the anaerobic chamber. Plates were then taken out. This was the initiation of the MA process. Caution was taken to ensure a strictly anaerobic condition maintained throughout the experiment. A single/tiny (< 1 mm) colony of each line was carefully picked and transferred onto a new plate. Then the new plates were put back into the anaerobic jar for incubation. The single cell bottleneck of the MA process occurred during every transfer.

The MA propagation was completed following 20 transfers, and four MA lines were lost during the MA process. A single colony on each plate was transferred into 5 mL anaerobic TRM medium in the anaerobic chamber. The liquid medium was supplemented with sulfur and $Na_2S{\cdot}9H_2O$. After incubation at 85°C for one day, stocks of each line were kept at -80°C. Genomic DNA of each survived MA line was extracted using the Magen Hipure Soil DNA Kit and sequenced using the same platform mentioned above. A sequencing coverage depth of ~433× with an average library fragment size of ~470 bp was obtained for each line.

*Generation time estimation with correction for cell death rate*

To estimate the generation time, a whole single colony was cut from 10 randomly selected MA lines. The selected 10 colonies each were moved into 5 mL anaerobic TRM medium supplemented with $Na_2S{\cdot}9H_2O$. After dilution and re-plating, live cell density ($d$) was measured with viable cell counts. The live and dead cell staining was done to correct the total cell density for each colony. Briefly, to obtain the sufficient cell density for staining, ten single colonies were cut from every MA line selected above. Live and dead bacterial staining kit (Yeasen Biotech Co.) was used in this study. The kit was tested to be effective in archaea. The cells were put into 350 μL anaerobic TRM medium supplemented with $Na_2S{\cdot}9H_2O$.

After centrifuge with 10,000 g for 10 min, cells were resuspended in 50 μL medium. Cell staining was done following the protocol of the kit. Fluorescence microscope (Nikon) was used to differentiate between live and dead cells. The ratio of live cells to total cells (r) was 0.942 (± 0.095) (Table S5). The number of cell divisions per transfer (*D*) was corrected by:

$$D = \log_2(\frac{d}{r})$$

where *d* is the live cell density and *r* is the ratio of live cells in total cells. The total number of generations that each MA line went through was the multiplication of average number of cell divisions per transfer and the total number of transfers. Since each MA line underwent 20 transfers with an average of 15.72 ± 1.76 cell divisions per transfer, there were a total of 314.4 ± 35.2 generations for each MA line.

*Mutation calling and mutation rate determination*

Raw reads were first processed by Trimmomatic 0.32 [4] to remove adaptors and trim low-quality bases. Then the paired-end reads of 96 MA lines were individually mapped to the *T. eurythermalis* A501 reference genome using two different mappers: BWA-mem [7] and NOVOALIGN v2.08.02 (www.novocraft.com). The resulting pileup files were converted to SAM format with SAMTOOLS [8].

The above mapping results were processed by Picard MarkDuplicates (http://broadinstitute.github.io/picard/) to remove duplicate reads which may arise during sample preparation like PCR duplication artifacts or derive from a single amplification cluster. Base quality score recalibration was performed to adjust quality score affected by systematic technical errors using BaseRecalibrator in GATK-4.0 [9]. Then base substitutions and small indels were called using HaplotypeCaller implemented in GATK-4.0 [9]. Variants were further filtered with standard parameters described by GATK Best Practices recommendations, except that the Phred-scaled quality score QUAL > 100 and RMS mapping quality MQ > 59 were set, which followed previous studies [9–12]. PCR primers were designed with Primer Premier 5.0 [13] to confirm the presence of mutations identified by the above bioinformatics method. Twenty base substitutions and nine indels were sampled from 11 lines and validated. These lines were chosen because two of these lines showed the highest base-substitution mutation rate and the remaining nine lines showed the longest indel mutations (Table S1). The average number of analyzable sites and the average coverage per site in the *T. eurythermalis* A501 MA lines were 2,123,047 (± 674) and 431 (± 57), respectively.

The base-substitution mutation rate per nucleotide site per cell division (μ) for each line was calculated according to the following equation:

$$\mu = \frac{m}{nG}$$

Where *m* is the number of observed base substitutions, *n* is the number of nucleotide sites analyzed, and *G* is the mean number of cell divisions estimated during the mutation accumulation process. Following a previous study [14], the total standard error of base-substitution mutation rate across all MA lines was calculated by:

$$SE_{pooled} = \frac{s}{\sqrt{N}}$$

where *s* is the standard deviation of the mutation rate across all lines, and *N* is the number of lines analyzed.

*The effective population size estimation for Thermococcus eurythermalis*

The effective population size ($N_e$) of a prokaryotic species was calculated following the equation $\pi_S = 2 \times N_e \times \mu$, where $\pi_S$ is the nucleotide diversity at silent (synonymous) sites among randomly sampled members of a species and μ is the unbiased spontaneous mutation rate. Microbial species commonly harbor genetically structured populations, which has a major influence on $\pi_S$ and thus $N_e$ estimation. It is therefore important to identify strains allowed for free recombination when calculating $N_e$ for a prokaryotic species [15]. The recently available program PopCOGenT [16] identifies members from a prokaryotic species constituting a panmictic population. The basic idea of PopCOGenT is that the recent homologous recombination erased the single nucleotide polymorphisms (SNPs) and led to identical regions between genomes, and therefore strains subjected with frequent recent gene transfers are expected to show an enrichment of identical genomic regions compared to accumulation of SNPs between genomes lacking recent transfer [16]. In practice, strains were connected via recent gene flow into a network, and a putative population was identified as a cluster, with within-cluster DNA transfer frequency much higher than that of between clusters. Only one strain within each clonal complex was kept, which is also important for $\pi_S$ estimation because an overuse of strains from a clonal complex is expected to underestimate $\pi_S$. Then the cluster containing the largest number of strains was chosen as the panmictic population for a given species. In the case of *T. eurythermalis,* all nine strains together form a panmictic population, but two strains were not used in the calculation because they were repetitive members of clonal complexes.

Next, the single-copy orthologous genes shared by all the seven *T. eurythermalis* genomes were identified by OrthoFinder 2.2.1 [17]. Amino acid sequences of each gene family were aligned with MAFFT v7.464 [18] and then imposed on nucleotide sequences. The number of synonymous substitution per synonymous site ($d_S$) for each possible gene pair in each gene family was computed with the YN00 program in PAML 4.9e [19]. The $\pi_S$ of each gene family was obtained by averaging all pairwise $d_s$ values, and then the median $\pi_S$ across all single-copy gene families together with μ were used to calculate the $N_e$. We used the median $\pi_S$ instead of the mean value, because loci showing unusually large $d_S$ as a result of allelic replacement via homologous recombination with divergent lineages are common in marine prokaryotic species [20], which are expected to bias the mean value but have a limited effect on the median value across gene loci. Given the small sample size of the available *T. eurythermalis* genomes, bootstrap resampling (with replacement, 10,000 pseudoreplicates) of the genomes were conducted to estimate standard deviation (Table S4) of $\pi_S$.

*Data synthesis*

To enable a comparative analysis of *T. eurythermalis* relative to other prokaryotic species, the available μ values of other 29 prokaryotic species determined with the MA/WGS technique were collected from the literature (Table S4). Among these, 20 species each had multiple isolates' genomes available from the NCBI Refseq database [21], and thus were used for $N_e$ calculation. The calculation of $N_e$ for these species followed the abovementioned procedure detailed for *T. eurythermalis*, which started with the identification of members constituting a panmictic population by PopCOGenT, followed by the calculation of $\pi_S$. The same bootstrap resampling analysis was performed to estimate the standard deviation of $\pi_S$ for another four species, each of which consists of less than 10 strains (Table S4). A few species have thousands of isolates' genomes available in Refseq (Table S4), which are not amenable for the PopCOGenT analysis. For these species, we started from the populations previously identified by ConSpeciFix [22, 23] and used these genomes as the input of PopCOGenT. The ConSpeciFix delineates populations based on homoplasious SNPs, which retains historical recombination signal and blurs the boundary of the ecological populations enriched with recent gene transfers [16]. In the case of the species *Ruegeria pomeroyi* DSS-3, a model heterotrophic marine bacterium with its mutation rate available [14], since closely related isolates has not been available, we turned to its closely related species *Epibacterium mobile* (previously known as *Ruegeria mobile*) with multiple isolates' genomes available.

Next, the pairwise linear relationship between $\mu$, $N_e$, and genome size across the prokaryotic species was initially assessed with the generalized linear model (GLM) implemented in *stats* package in R v4.0.2 [24]. The Bonferonni adjusted outlier test was performed with *outlierTest* function in *car* package [25]. A data point with Bonferroni *p*-value smaller than 0.05 would be identified as the outlier. For $\mu$ versus genome size, all 30 species were used. In the case of $N_e$ versus $\mu$ and $N_e$ versus genome size, only the 21 species each containing multiple strains' genomes were used. To test whether there was a phylogenetic signal of these traits, the Pagel's λ [26] was estimated using the *pgls* function of the *caper* package [27] which took the phylogeny of 30 species or the phylogeny of 21 species as an input. The species phylogeny was approximated by the 16S rRNA gene tree constructed using IQ-TREE 2.0 [28] with ModelFinder [29] which assigns the best substitution model and with 1,000 ultrafast bootstrap replicates. The value of λ ranges from 0 to 1, with 0 indicating no phylogenetic signal and 1 indicating a strong phylogenetic signal due to Brownian motion. The *p* values for the lower and upper bounds represent whether the λ is significantly different from 0 and 1, respectively. The results of this test indicate that there was an intermediate phylogenetic signal for the relationship of $N_e$ versus μ ($\lambda = 0.81$, lower bound $p = 0.29$, upper bound $p = 0.06$), but not for that of $N_e$ versus genome size and μ versus genome size (in both cases, $\lambda = 0$, lower bound $p = 1$, upper bound $p < 0.001$). To control for the phylogenetic effect on the correlations of the traits, the pairwise linear relationship between μ, $N_e$, and genome size was further assessed with the phylogenetic generalized least square (PGLS) regression implemented in the *caper* package [27] in R v4.0.2 [24]. The PGLS and GLM regression lines were largely overlapped for $N_e$ versus genome size and μ versus genome size (Fig. 2BC). This is because no phylogenetic signal was detected in these relationships. A data point was identified as an outlier in the PGLS result if the associated absolute value of studentized residual is greater than three [30, 31].
